# How do visual and conceptual factors predict the composition of typical scene drawings?

**DOI:** 10.1101/2025.09.15.676247

**Authors:** Gongting Wang, Ilker Duymaz, Matthew J. Foxwell, Micha Engeser, David Pitcher, Radoslaw M. Cichy, Daniel Kaiser

## Abstract

Imagine you are asked to draw a typical bedroom, what would you put on paper? Your choice of objects is likely to depend on visual occurrence statistics (i.e., the objects present in previously encountered bedrooms) and semantic relations between objects and scenes (i.e., the semantic relationship between the bedroom and its constituent objects). To investigate how these two factors contribute to the composition of typical scene drawings, we analyzed 1,192 drawings of six indoor scene categories, obtained from 303 participants. For each object featured in the drawings, we estimated its visual occurrence frequency from the ADE20K dataset of annotated scene images, and its semantic relatedness to the scene concept from a word2vec language processing model. Across all scenes of a given category, generalized linear models revealed that visual and conceptual factors both predicted the likelihood of an object featuring in the scene drawings, with a combined model outperforming both single-factor models. We further computed the visual and semantic specificity of objects for a given scene, that is, how diagnostic an object is for the scene. Object specificity offered only weak predictive power when predicting the selection of objects, yet even infrequently drawn objects remained diagnostic of their scenes. Taken together, we show that visual and conceptual factors jointly shape the composition of typical scene drawings. By releasing a large dataset of typical scene drawings alongside this work, we further provide a starting point for future studies exploring other critical properties of human drawings.

## Introduction

Drawing is a fundamental form of human communication. Humans have employed drawings to communicate ideas, express emotions and share experiences for at least 40,000 years (Aubert et al., 2014; Hoffmann et al., 2018). In modern society, drawings continue to serve diverse functions. For instance, designers use drawings to visualize ideas (Goldschmidt., 2014), artists use drawings to externalize imagination (Fish & Scrivener., 1990), clinicians use drawings to diagnose and classify neurological disorders (Agrell., 1998; Wechsler., 2009). During many everyday activities, people draw to alleviate boredom or to enhance concentration (Andrade., 2010). Contrasting this prevalence of drawings, we know relatively little about how people decide which elements to include when composing a drawing. This question is particularly evident when drawings of complex scenes are considered: which objects do people draw to convey a certain scene, say a living room? A better understanding of the composition of such drawings may shed a new light on how people translate perceptual and memory representations into visible outputs.

Such insights are also crucial for researchers in cognitive science and neuroscience that use drawing as a tool to characterize the nature of internal representations (Engeser et al., 2025; Fan et al., 2023; Roberts & Wammes, 2021). For instance, in development research, drawings are used to investigate the developmental trajectory of visual object representations (Karmiloff-Smith., 1990; Long et al., 2019; Long et al., 2024). In memory research, successes and failures in memory can be captured by how well human drawings during recall align with or deviate from the studied materials (Metzger., 1936; Bainbridge et al., 2019; Bainbridge & Baker., 2020; Fan et al., 2023). Further, in perception research, drawings provide descriptions of participants’ world models, which can in turn be used to predict perception (Engeser et al., 2025; Morgan et al., 2019; Wang et al., 2024, 2025). A better understanding of how people compose their drawings could inform the potential and limitations of studies using drawing as a methodological tool.

To understand how people compose drawings of natural scenes, we analyzed more than 1,000 drawings of six scene categories from more than 300 participants. All participants were asked to draw a typical instance of a scene category (e.g., a typical living room), after briefly thinking about the scene contents and then drawing within a relatively liberal time constraint (Wang et al., 2024, 2025). The full set of scene drawings, the *Room Drawings Dataset*, is released alongside this publication (see Materials and Methods), providing a rich benchmark dataset of evaluating diverse aspects of drawing composition in future studies.

In the current study, we used the dataset to ask how well the object composition in drawings from each scene category could be predicted by two complementary factors: (i) *visual occurrence statistics*, that is, the frequency with which the individual objects are encountered in a given scene category in the real world, and (ii) *semantic similarity*, that is, the semantic relatedness between the scene concept and the individual object concepts.

First, people’s decisions about which objects are drawn for a given category should predict on how frequently the objects are commonly found in scenes of that category. In bathrooms, we much more often see a sink than a table, so we should draw sinks more often than tables. Most scenes are reliably associated with such prominent scene-and space-defining objects (Bar, 2004; Bar & Aminoff, 2003; Oliva & Torralba, 2007; Võ et al., 2019). Here, we quantified such *visual occurrence statistics* by assessing how frequently individual objects appear within images real-world scene categories. Specifically, we used the annotated ADE20K scene database (Zhou et al., 2017; Zhou et al., 2018) and computed for a set of scene categories how often individual objects featured in images of that category (e.g., how often is a sink found in images of bathrooms).

Second, the object composition of a drawing may not just be shaped by visual occurrence statistics. It may also depend on information stored in conceptual representations, which do not necessarily mirror unfiltered visual experience. During drawing, people need to recall relevant objects from long-term memory. Such long-term memory for real-world objects can be stored in structured conceptual spaces (Brady et al., 2008; Konkle et al., 2010), and retrieval from such conceptual spaces may depend on the semantic similarity between a scene concept and the candidate object concepts. Here, we quantified such *semantic similarity* by assessing the similarity of object and scene concepts in a large language model (word2vec; Mikolov et al., 2013). Specifically, we computed the cosine similarity between a set of scene category concepts and individual object concepts in this model (e.g., how similar is the concept “sink” to the concept “bathroom”).

These two predictors enabled us to test how visual occurrence statistics and semantic relatedness contribute to the object composition in a set of more than 1,000 scene drawings spanning six categories.

## Materials and methods

### Participants

A total of 303 participants (26.14±4.78 years±SD, 100/201/2 male/female/other) provided scene drawings. Of these, 101 participants (25.20 ±4.07 years±SD, 24/77 male/female) were tested online in the UK (recruited at the University of York), and 202 participants (26.62±5.04 years, 76/124/2 male/female/other) were tested in a laboratory setting in Germany (recruited at Freie Universität Berlin and Justus-Liebig-Universität Giessen). Procedures were approved by the ethics committees of the Department of Psychology, University of York, the Department of Education and Psychology, Freie Universität Berlin, and the ethics committee of the Justus-Liebig-Universität Gießen, respectively, and adhered to the Declaration of Helsinki.

### Drawing sessions

The drawings used here were produced in drawing sessions across multiple experiments, including two published studies (Wang et al., 2024, 2025) and other still unpublished work. Drawing sessions were conducted either online or in-person in a laboratory setting. Online sessions were conducted via Skype. Here, participants provided their drawings using a pencil, eraser, and ruler on A4 paper. For each drawing, participants had 1 minute to plan and 3 minutes 30 seconds to complete the drawing. Participants drew typical versions of three scene categories (bedroom, kitchen, living room). In the lab-based drawings sessions, participants provided their drawings using an Apple Pencil on an Apple iPad with the Sketchbook App. Each participant was given 30 seconds to plan and 4 minutes to complete each drawing. A group of participants (N=85) drew six typical scene categories (bathroom, bedroom, café, kitchen, living room, office), while another group (N=115) drew four categories (bathroom, bedroom, kitchen, living room). In all sessions, participants were instructed to draw the most typical representation of each room (Figure 1), rather than their own rooms or an aesthetically appealing version of the room. Participants were instructed to draw into a pre-defined perspective grid, which they drew themselves according to the experimenter’s instructions (online sessions) or which was present on the iPad screen from the outset (lab-based sessions). Further details on the drawing sessions are available in our previous studies (Wang et al., 2024, 2025).

**Figure 1.**
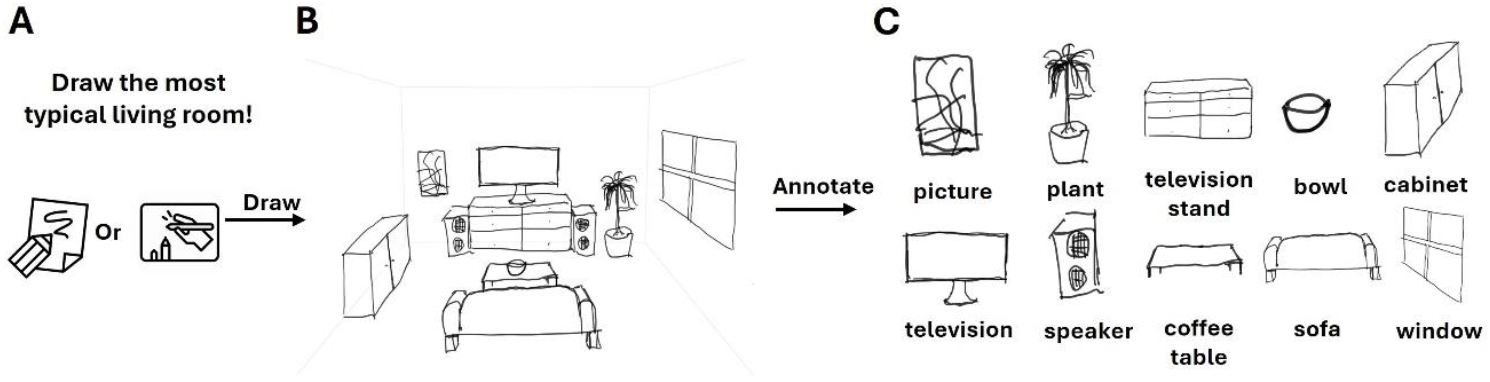
Overview of the drawing sessions and annotation procedure. **(A)** Participants draw typical versions of indoor scene categories on paper or an iPad. **(B)** An example drawing of a living room. Note that a common perspective was enforced by a perspective grid shown to the participants from the outset. **(C)** We manually annotated all individual objects in the drawings. Each object instance was only counted once (i.e., objects were coded as present or absent).

### Object annotation

For each drawing, all depicted objects were manually annotated. Overall, typical bathroom drawings consisted of an average of 6.3 different objects (SD = 1.6, range = [4, 12]; multiple instances of the same object were not counted), with the three most frequent objects sink (100%), toilet (96%), and shower (86%). Bedrooms consisted of an average of 8.3 objects (SD=2.1, range = [4, 13]), with the three most frequent objects bed (100%), pillow (93%), and window (74%). Cafés consisted of an average of 7.4 objects (SD=3.1, range= [3, 19]), with the three most frequent objects table (99%), chair (97%), and counter (86%). Kitchens consisted of an average of 8.7 objects (SD= 3.4, range = [4,17]), with the three most frequent objects cupboard (99%), stove (98%), and sink (90%). Living rooms consisted of an average of 7.8 objects (SD=2.5, range = [4,14]), with the three most frequent objects sofa (100%), television (89%) and television stand (77%). Offices consisted of an average of 7.3 objects (SD =3.1, range = [4, 15]), with the three most frequent objects table (100%), chair (94%), and window (75%). Full annotation details are provided in the supplementary materials. From these annotations, we computed the occurrence frequency of each object within its respective scene category.

### Quantifying visual occurrence statistics and semantic relatedness

To quantify visual occurrence statistics across the exemplars of each scene category, we determined the occurrence frequencies of the annotated objects in each scene category by referencing the ADE20K dataset (Zhou et al., 2017; Zhou et al., 2018). Specifically, we queried the database for each object annotated in the drawings and the corresponding scene category and calculated each object’s frequency of occurrence across all the scenes for each category. To quantify semantic relatedness between object and scene concepts, we computed the similarity between the word representing each object annotated in the drawings (e.g., “shower” in English, “Dusche” in German) and the corresponding scene concepts (e.g., “bathroom” in English, “Badezimmer” in German) using a word2vec model. We used word vectors pre-trained on German for the German participants and on English for the UK participants, and supplied words in each group’s native language. Both training resources came from the Common Crawl and Wikipedia corpora using fastText (Grave et al., 2018). By calculating the cosine similarity between the object and scene concept, we quantified how strongly each object is semantically related to the scene category it appeared in.

### Modelling object drawing frequencies

To examine the relationship between object drawing frequency, visual occurrence statistics, and semantic relatedness, we fitted generalized linear models to predict object drawing frequencies using (i) occurrence statistics only, (ii) semantic relatedness only, and (iii) both predictors together. As the dependent variable represented a bounded proportion (i.e., % of drawings that contained the object), we chose a Beta-binomial regression approach (i.e., a generalized linear model with a Beta function as the link function, Ferrasi & Cribari-Neto, 2004). To assess the overall effect of visual experience and conceptual knowledge on object occurrence frequency across all categories, we fitted a generalized linear mixed-effects model that included category as a random effect as well as visual experience and/or conceptual knowledge as fixed effects. We then assessed whether a combined model better explained drawing frequencies better than occurrence statistics or semantic relatedness along. To assess the stability of the results across categories, we further fitted the same model individually for every scene category.

We explored the composition of the drawings further by asking how the specificity of an object for a given scene category (e.g., a stove is highly specific for a kitchen, as it almost exclusively appears there, but a window is not) predicts drawing frequency. To quantify specificity, we calculated (i) the scene-specificity of an object in visual occurrence statistics, defined as the normalized difference between an object's frequency in its corresponding scene category and its average frequency across the other scene categories, the (ii) the scene-specificity of an object in semantic relatedness, defined as the normalized difference between an object concept's cosine similarity to its corresponding scene concept and its average cosine similarity to the other scene concepts in the language model. Specificity was computed using the following formula:

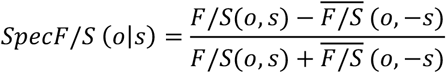

F: object frequency, S: semantic relatedness, o: object, s: corresponding scene, -s: the other scenes.

Then, we fitted mixed effects models to predict object frequency in drawings from visual and semantic specificity. We further asked whether infrequently drawn objects are still diagnostic for their respective scene category (and if so, to which extent) or whether they mainly constitute generic “filler” objects that are equally appropriate for our range of categories (like windows or bins). To this end, we ordered all objects by their drawing frequency and binned them into six ranges: objects featured in 0-10%, 10-20%, 20-30%, 30-40%, 40-70%, or 70-100% of scenes. We varied bin sizes across the frequency range as there were more objects that appeared relatively infrequently. We then assessed whether objects across frequency bins were consistently diagnostic for the scene category.

The ADE20K annotations analysis and the word2vec analysis were conducted in Python. All further statistical analyses were conducted in R.

## Data, material and code availability

Data, and code are accessible on the Open Science Framework (OSF), available at https://osf.io/p2fa6/. We further release all drawings, together with participant information and annotations, in the *Room Drawings Dataset*, available at: https://osf.io/byu24/.

## Results

We first visualized how often different objects were drawn for each scene category. We generated word clouds (Heimerl et al., 2014) for each category, in which font size reflects how frequently each object was drawn (Figure 2). These descriptive data show that each scene featured prominent and diagnostic objects that were drawn across many instances of the scene (e.g., a bed in the bedroom).

**Figure 2.**
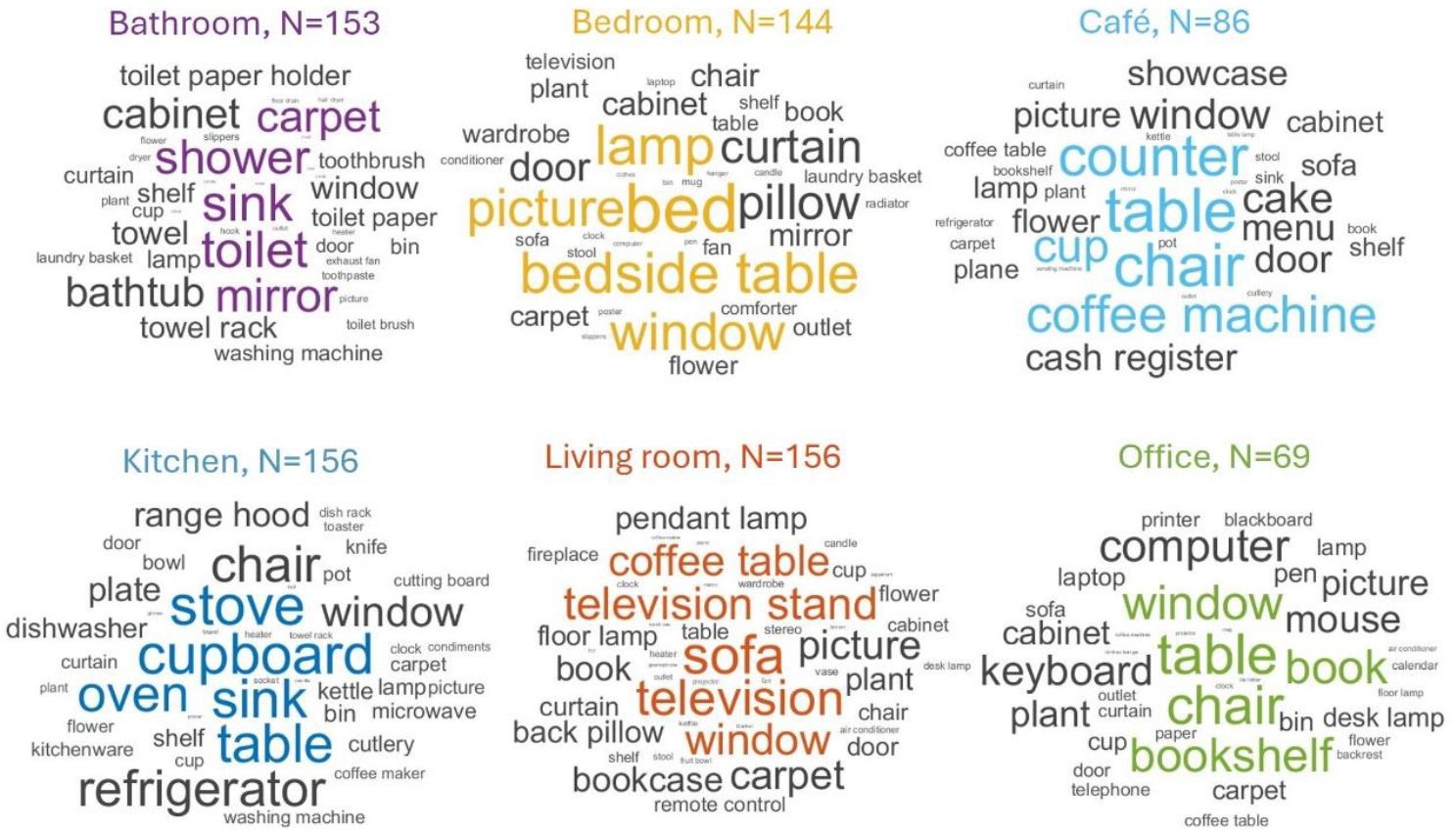
Word clouds representing the drawing frequency of objects across scene categories. The size of the words represents how frequently the object was featured in the drawings.

To evaluate how visual occurrence statistics and semantic relatedness contribute to the composition of scene drawings, we fitted three mixed-effects models that predicted object drawing frequency from either (i) only visual occurrence statistics, (ii) only semantic relatedness, or (iii) both factors combined. Each model additionally featured category as a random effect. Model estimates and fit indices are reported in Table 1 (“Full Model”). A comparison of Akaike Information Criterion (AIC) and Bayesian Information Criteria (BIC) revealed that the combined model provided the best fit (AIC = -679.50, BIC = -659.98), with a conditional R^2^ of 0.81. These results suggest that visual occurrence statistics are a good predictor of object drawing frequencies in scene drawings. Yet, semantic similarity between scene and object concepts explains additional variance in the drawing composition that is not captured by visual statistics.

**Table 1.** Regression weights and goodness-of-fit statistics for the generalized linear models.

| Scene Category | Visual | Semantic | AIC | BIC | R<sup>2</sup> |
| --- | --- | --- | --- | --- | --- |
| Full Model | 4.78*** | / | -645.65 | -630.04 | 0.77 |
|  | / | 6.59*** | -436.85 | -421.24 | 0.54 |
|  | 4.33*** | 3.21*** | -679.50 | -659.98 | 0.81 |
| Bathroom | 4.39*** | 5.99*** | -92.89 | -86.12 | 0.67 |
| Bedroom | 4.56*** | 2.10* | -156.11 | -146.39 | 0.64 |
| Café | 6.03*** | 10.37*** | -56.86 | -50.52 | 0.65 |
| Kitchen | 4.36*** | 2.44* | -138.78 | -129.10 | 0.61 |
| Living room | 3.63*** | 3.30*** | -158.63 | -148.86 | 0.51 |
| Office | 6.69*** | 2.69* | -89.46 | -82.91 | 0.74 |

Moreover, the random effect of category was significant (p<0.001). We thus examined whether predictor contributions varied across scene categories, we then fitted separate generalized linear models with both predictors in each category (Figure 3). In all cases, both visual occurrence statistics and semantic relatedness remained significant, although their relative contributions varied across categories (Table1). The generalized linear models also predicted object drawing frequency when trained on all but one category and tested on the remaining category (see Supplementary Information), suggesting an overall similar influence of visual occurrence statistics and semantic relatedness across categories. Finally, the complementary contribution of visual and semantic factors was corroborated by a simpler partial correlation analysis, where partialing out visual occurrence statistics still yielded significant correlations between semantic similarity and object drawing frequencies, and vice versa (see Supplementary Materials).

**Figure 3.**
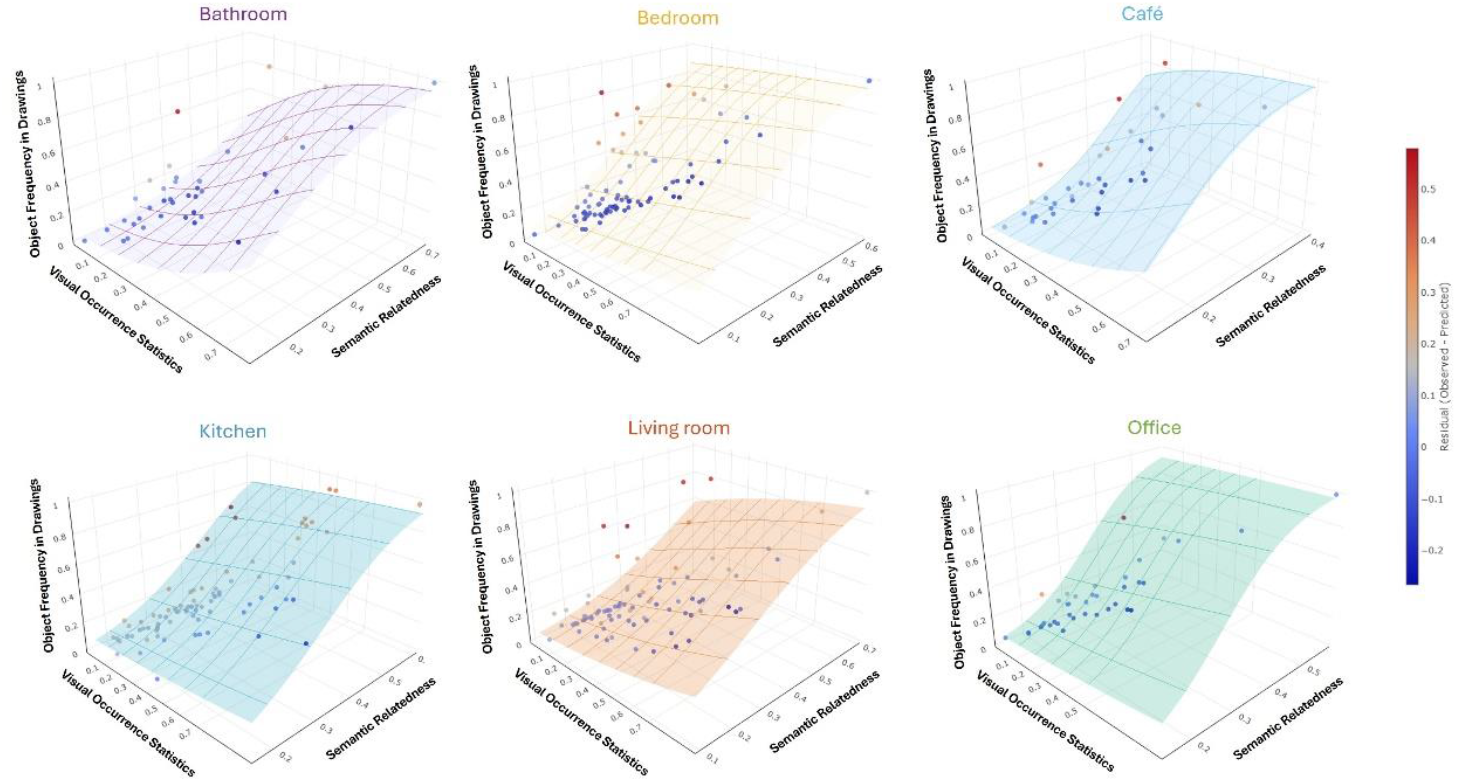
Visualization of generalized linear model fits when predicting object drawing frequencies from visual occurrence statistics and semantic relatedness, separately for the six categories. In all categories, both predictors yielded coefficients significantly greater than zero. Except for the café category, the coefficient for visual occurrence statistics exceeds that for semantic relatedness, indicating a stronger influence of visual experience on drawing composition.

While the above analyses show that both visual and semantic factors contribute to whether objects are features in a drawing, it remains unclear whether participants select objects primarily based on absolute frequencies (i.e., how often does a chair appear in a living room) or based on relative frequencies (i.e., how much more often does a char appear in a living room compared to other scenes). Thus, we fitted mixed-effects GLMs that predicted object drawing frequency from visual and semantic specificity, again comparing single-factor to combined-factor models (Table 2). The results showed that the combined-factor model including both specificity predictors did not outperform the model that predicted drawing frequency from scene-specificity in visual occurrence statistics alone. This suggests that when selecting objects based on specificity, visual specificity (i.e., whether an object is more often seen in one room compared to other rooms) trumps semantic specificity (i.e., whether an object is semantically associated more strongly with one category than with the others).

**Table 2.** Regression weights and goodness-of-fit statistics for generalized linear models with specificity predictors.

| Scene Category | Visual | Semantic | AIC | BIC | R<sup>2</sup> |
| --- | --- | --- | --- | --- | --- |
| Full Model | 0.55<sup>***</sup> | / | -343.73 | -328.12 | 0.13 |
|  | / | 1.16<sup>**</sup> | -328.38 | -312.77 | 0.05 |
|  | 0.56<sup>***</sup> | -0.05 | -341.73 | -322.22 | 0.13 |

However, it is worth noting that overall model performance in this analysis was relatively low, as reflected in the low Rsss values. This suggests that specificity alone might not drive object selection in drawings. Rather than selecting objects because they are highly specific to a scene, participants may instead choose objects based on how frequently they appear in a given contexts regardless of how often they appear elsewhere. This raises another question: for infrequently drawn objects, are people simply selecting objects that are broadly associated with many scenes and thus go well with everything? Or were these infrequent objects still scene-diagnostic? To address this, we utilized our specificity measure to examine whether even infrequently drawn objects were still diagnostic for the scene categories they were drawn in. As expected, frequently drawn objects (those drawn in at least 40% of scenes) showed clear specificity both in visual occurrence statistics (all t > 4, all p < 0.0001) and semantic specificity (all t > 3, all p < 0.01). Critically, our analysis showed that even infrequent objects (those drawn in less than 40% of scenes) retained significant specificity across both visual (all t > 2, all p < 0.05) and semantic measures (all t > 2, all p < 0.05; Figure 4A, 4B), suggesting that infrequently included objects are still specifically associated with the target scene category (e.g., a hair dryer as a low-frequency object with high specificity for a bathroom), rather than simply being generic or universally compatible.

**Figure 4.**
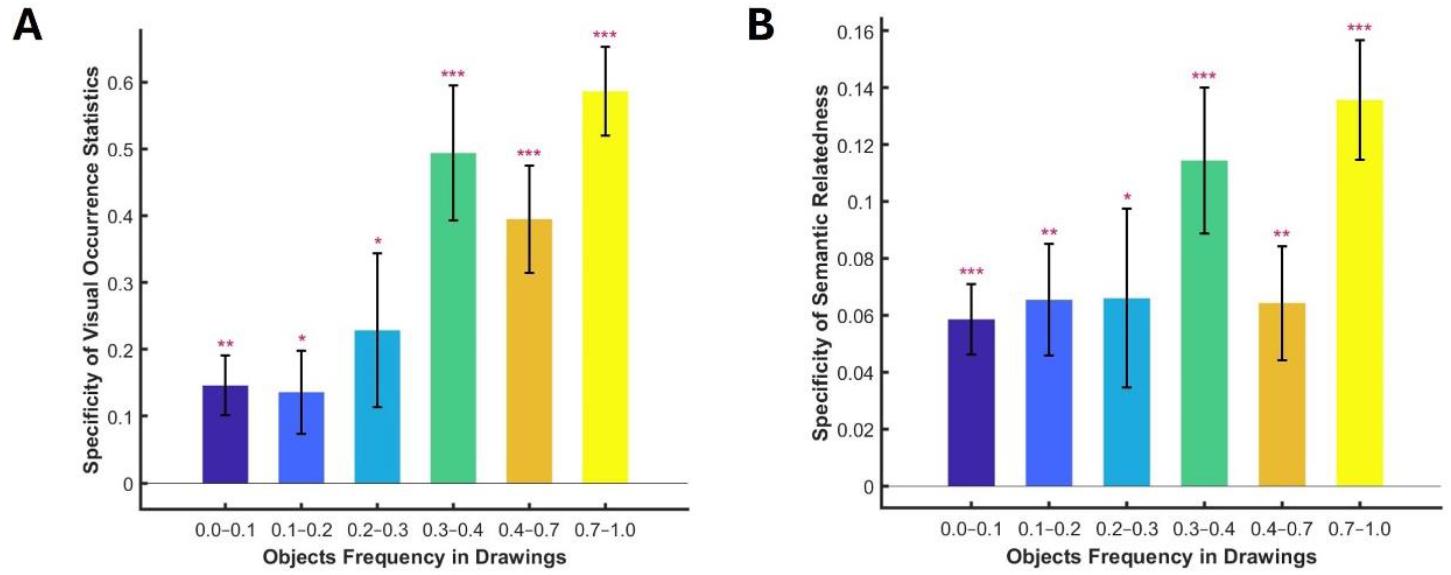
Assessing visual and semantic specificity of frequent and rare objects in scene drawings. Both (A) visual specificity and (B) semantic specificity are significantly positive across drawing frequency bins, suggesting that even rarely drawn objects are, on average, scene-diagnostic. Error margins represent s.e.m. ^*^p<0.05 ^**^p<0.01^***^p<0.001

## Discussion

This study examined how visual occurrence statistics and semantic relatedness determine which objects are drawn when participants compose drawings of typical real-world scenes. Across six scene categories, we demonstrate that both factors significantly predicted how often objects were included in the drawings, and a combined model explained more variance than either predictor alone. Nonetheless, visual occurrence statistics consistently emerged as the stronger predictor. How often objects appear in scenes of a certain category and how strongly they are related to the scene concepts predicted object drawings frequencies better than the scene-specificity of the objects (i.e., whether an object is more often found in or more strongly to the drawn scene category than to the other scene categories). Yet, even objects that were drawn infrequently were, on average, diagnostic for the scene category they were included in.

Producing a typical scene drawing necessitates the retrieval of relevant objects from long-term memory. Based on the classical schema theory (Biederman et al., 1982; Boyce et al., 1989), participants initially active a semantic representation of the scene category from long-term memory. They then use prediction of expected semantic associations between the objects and scenes to guide visual information gathering (Bar, 2004; Oliva & Torralba, 2007; Leroy et al., 2020). These scene representations constrain the candidate objects that belong in the corresponding scene. The subsequent retrieval of objects is likely constrained by the structure of the scene, such as the typical spatial distributions of objects (Bar, 2004; Kaiser et al., 2019; Kaiser et al., 2015; Võ et al., 2019) and the spatial layout of whole scenes (Kaiser & Cichy, 2021). Critically, detailed representations of objects within scenes rely heavily on visual long-term memory (Brady et al., 2008), which is shaped by everyday visual statistical learning (Stansbury et al., 2013). Through repeated exposure to visual occurrence statistics in daily life, participants implicitly learn the typical composition and spatial arrangement of objects, enabling them to accurately predict and reconstruct detailed object information during the drawing process. Thus, both semantic and visual formats jointly contribute to the retrieval and representation of typical scenes. Interestingly, the observed dominance of visual occurrence statistics in predicting object frequency in drawings may reflect the fundamental role of real-world visual experience in shaping scene representations stored in long-term memory. Specifically, repeated visual encounters with objects in particular scene contexts likely strengthen their internal representations, making these frequently encountered objects more easily retrievable and visually detailed during reconstruction (Brady, Konkle, & Alvarez, 2009; Torralba et al., 2016).

On the other hand, the relatively weaker performance of our semantic predictor might partly stem from the hubness problem in word2vec-based semantic measurement (Schnabel et al., 2015). Specifically, when words are projected into high-dimensional vector spaces, “hubs” appearing as nearest neighbors to a disproportionately large number of other points (Radovanovic et al., 2010). For example, “kitchen” becomes a hub that attracts objects to similar distances, which caused a narrow similarity range. This problem might compress the distribution of semantic associations, causing many scene-object pairs to tightly cluster within a restricted similarity band. Therefore, the low variance likely limits the measure's ability to explain frequency values. Future studies could apply models that yield more variability and thus diagnostic power, such as BERT-based (Devlin et al., 2019) and CLIP-based (Radford et al., 2021) measurements.

### nau mattias

Furthermore, scene representations stored in memory are rooted in differences in participants' visual diets and may thus differ substantially between individuals (Engeser et al., 2025). In this study, we utilized object frequencies derived from the ADE20K image database and semantic associations from word2vec model as proxies for real-world visual experience and conceptual knowledge. Despite essentially ignoring all inter-individual variance, this approach still predicted drawing content, highlighting the potential of this method to assess internal visual representations. Future research could incorporate more individualized measures, such as individual photographic exposure logs, personal digital archives, or models trained on bespoke cultural and linguistic backgrounds. Such refinements might better characterize visual and semantic contributions on the individual or group level, thereby yielding more accurate predictions from visual and semantic factors.

Interestingly, the specificity of an object for a given scene category (i.e., whether an object was by itself diagnostic for the scene category) was a relatively weak predictor of object drawing frequency. This suggests that participants adopted a task-specific mindset in which they mainly focused on absolute object occurrence statistics that were best suited to maximize scene typicality (Wiesmann & Võ., 2023). If our task shifted from “draw a typical bedroom” to “draw the most diagnostic bedroom,” object selection should pivot towards a greater weighting of specificity, increasing the contribution of items that are perhaps rarer but more exclusive to the category. It is also worth noting that our specificity measure was only computed relative to the five alternative categories in the current set. Future work could compute the scene-specificity of objects in relation to a broader set of reference categories to more accurately gauge specificity.

An additional contribution of this work is the public release of a large, object-annotated database of human typical scene drawings, the *Room Drawings Dataset*, which containing more than one thousand drawings across six categories (see Materials and Methods). Because each drawing comes with category and object labels and participant information, the dataset supports a wide range of secondary analyses, for instance, training and testing computational models on the drawings; comparing visual vs. semantic predictors across populations; benchmarking typical scene construction in clinical or developmental cohorts; and linking typical drawing content to scene-selective neural responses. Furthermore, the dataset allows for easily appending new drawings, new scene categories, additional object codes, or supplemental metadata. We hope this resource will serve as a useful resource for modeling scene perception and cognition.

In sum, our findings reveal visual occurrence statistics and semantic relatedness jointly predict the composition of typical scene drawings. Visual frequency exerts the stronger influence on which core objects are rendered, and even infrequent drawn objects still contributing to category identity. These insights provide a new behavioral window onto the internal representations of the world that support human scene understanding across individuals.

## Supplementary Materials

### Partial correlation analysis

To disentangle the contributions of visual occurrence statistics and semantic relatedness, we also conducted partial correlation analyses within each scene category. Controlling for semantic relatedness, object frequency in drawings remained strongly correlated with visual occurrence statistics in every category: bathroom: r=0.66, p<0.001; bedroom: r=0.61, p<0.001; café: r=0.74, p<0.001; kitchen: r=0.72, p<0.001; living room: r=0.60, p<0.001; office: r=0.87, p<0.001; Conversely, when controlling for visual occurrence statistics, object frequency remained significantly correlated with semantic relatedness in most categories: bathroom: r=0.51, p=0.001; bedroom: r=0.24, p=0.03; café: r=0.60, p <0.001; living room: r=0.33, p=0.002; kitchen: r=0.24, p=0.03. but not office: r=0.07, p=0.71. These results confirm that both predictors uniquely account for variance in object drawing frequency, with visual occurrence statistics offering particularly robust predictions.

### Cross-validation analysis

To evaluate the model’s ability to generalize across scene categories, we conducted a leave-one-category-out cross validation using a beta-binomial regression with visual occurrence statistics, semantic relatedness and both factors as predictors, separately. In each iteration, the model was trained on five categories and tested on the sixth. Predictive accuracy was assessed by the coefficient of determination,

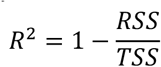

where RSS is the residual sum of squares and TSS is the total sum of squares. The detailed R^2^ was listed in Table S.

**Table S.** R^2^ for models trained on visual only, semantic only and combined factors.

| $R^2$ \ Models | Visual only | Semantic only | Combined |
| --- | --- | --- | --- |
| Bathroom | 0.56 | 0.25 | 0.67 |
| Bedroom | 0.47 | 0.25 | 0.48 |
| Café | 0.32 | 0.14 | 0.43 |
| Living room | 0.22 | 0.02 | 0.35 |
| Kitchen | 0.57 | 0.07 | 0.56 |
| Office | 0.63 | 0.22 | 0.67 |
| Averaged | 0.46 | 0.16 | 0.53 |

## Acknowledgements

This work was supported by the Deutsche Forschungsgemeinschaft (DFG) under Germany’s Excellence Strategy (EXC 3066/1 “The Adaptive Mind”, project no. 533717223). It was further supported by a European Research Council (ERC) Starting Grant (PEP, ERC-2022-STG 101076057). Views and opinions expressed are those of the authors only and do not necessarily reflect those of the European Union or the European Research Council. Neither the European Union nor the granting authority can be held responsible for them. The authors thank Daniela Marinova, Nasibeh Babaei and Pietra Pacheco Alves for their support in collecting the drawings.

